# NucPosDB: a database of nucleosome positioning *in vivo* and nucleosomics of cell-free DNA

**DOI:** 10.1101/2021.11.24.469884

**Authors:** Mariya Shtumpf, Kristan V. Piroeva, Shivam P. Agrawal, Divya R. Jacob, Vladimir B. Teif

**Author notes:** Correspondence should be addressed to Vladimir B Teif.

## Abstract

Nucleosome positioning is involved in many gene regulatory processes happening in the cell and it may change as cells differentiate or respond to the changing microenvironment in a healthy or diseased organism. One important implication of nucleosome positioning in clinical epigenetics is its use in the “nucleosomics” analysis of cell-free DNA (cfDNA) for the purpose of patient diagnostics in liquid biopsies. The rationale for this is that the apoptotic nucleases that digest chromatin of the dying cells mostly cut DNA between nucleosomes. Thus, the short pieces of DNA in body fluids reflect the positions of nucleosomes in the cells of origin. Here we report a systematic nucleosomics database – NucPosDB, curating published nucleosome positioning datasets *in vivo* as well as datasets of sequenced cell-free DNA (cfDNA) that reflect nucleosome positioning *in situ* in the cells of origin. Users can select subsets of the database by a number of criteria and then obtain raw or processed data. NucPosDB also reports the originally determined regions with stable nucleosome occupancy across several individuals with a given condition. An additional section provides a catalogue of computational tools for the analysis of nucleosome positioning or cfDNA experiments and theoretical algorithms for the prediction of nucleosome positioning from DNA sequence. We provide an overview of the field, describe the structure of the database in this context and demonstrate data variability using examples of different medical conditions. NucPosDB is useful both for analysis of fundamental gene regulation processes and training computational models for patient diagnostics based on cfDNA. The database currently curates ∼400 publications on nucleosome positioning in cell lines and *in situ* as well as cfDNA from >10,000 patients and healthy volunteers. For open-access cfDNA datasets as well as key MNase-seq datasets in human cells, NucPosDB allows downloading processed mapped data in addition to the stable-nucleosome regions. NucPosDB is available at https://generegulation.org/nucposdb/.

## Background

Genomic nucleosome positions are non-random and unique for each cell, reflecting many biological processes that require the access of regulatory molecules to the DNA (e.g. reviewed in (Clarkson et al. 2019; Baldi et al. 2020; Parmar and Padinhateeri 2020)). Previously, we assembled a comprehensive collection of experimental datasets of nucleosome positioning across many organisms and cell lines as well as software tools for the analysis and prediction of nucleosome positioning (Teif 2016). After the initial focus on nucleosome positioning in organisms such as Yeast (Yuan et al. 2005; Ioshikhes et al. 2006; Segal et al. 2006) many studies focused on human cells (Schones et al. 2008; Valouev et al. 2011; Gaffney et al. 2012; Kundaje et al. 2012; Diermeier et al. 2014; Ho et al. 2014; Teif et al. 2017; Mallm et al. 2019). Furthermore, more recently the field has moved towards clinical applications of nucleosome positioning to cell-free DNA (cfDNA), as will be explained below. There is a strong need for an integrative database that connects both fundamental and clinically focused “nucleosomics”. Here we report a systematic database, called *NucPosDB*, which integrates classical nucleosome positioning studies with a new direction of nucleosome positioning landscapes reconstructed from cfDNA from human patients.

The shift of the focus of the research from fundamental roles of nucleosome positioning in gene regulation to patient diagnostics is happening due to the fact that nucleosome positioning can provide a valuable diagnostic marker offering unique features not available in other clinical tests. There are two main arguments for this. Firstly, the timescale of the change of nucleosome positioning landscape is comparable to the timing of gene activation or the cell cycle (Schones et al. 2008; Teif et al. 2012) which is between the quick changes of concentrations of disease-related small molecules and the much slower changes reflected by DNA mutations or aberrant methylation happening in cancer (Dawson and Kouzarides 2012; Pich et al. 2018; Li and Luscombe 2020) (Figure 1A). Thus, differences in nucleosome positioning can be in principle suitable for monitoring a patient’s response to therapy in this intermediate time range. While very informative, determining genome-wide nucleosome positioning maps in tumour tissues of cancer patients would be an expensive and invasive procedure. Here, the second argument comes into play: luckily, nucleosome positioning in tissues is directly reflected in cfDNA circulating in blood and other body liquids. This is because nucleases, which shred the chromatin of dying cells to form what later becomes cfDNA, preferentially cut the DNA between nucleosomes (Chandrananda et al. 2015; Kustanovich et al. 2019; Serpas et al. 2019; Han et al. 2020; Heitzer et al. 2020) (Figure 1B). Since the half-life of cfDNA in blood is about 15 minutes (Volik et al. 2016), cfDNA extracted at any given time point represents a very recent snapshot of nucleosome positioning in the cells of origin.

**Figure 1.**
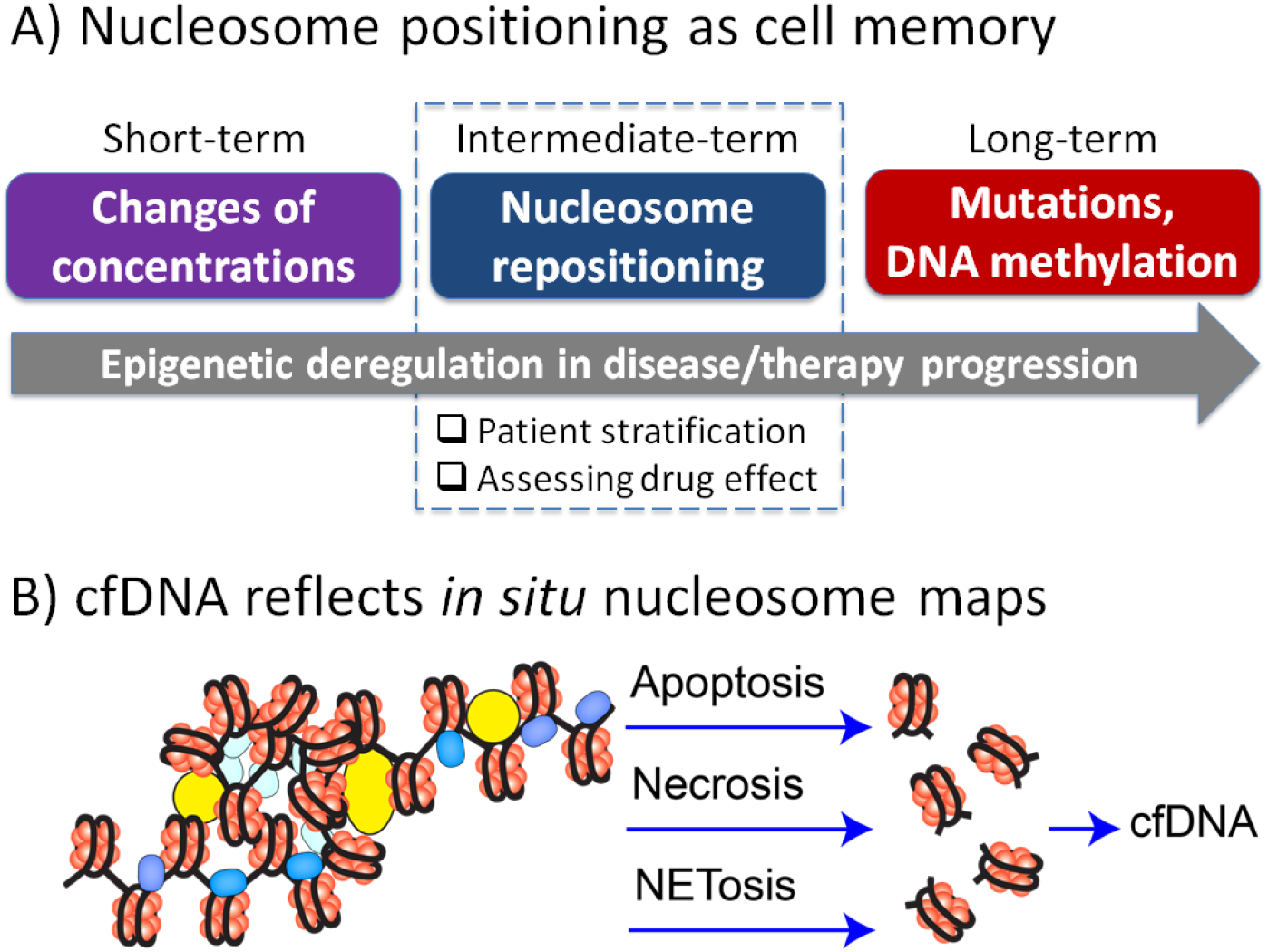
Motivation for the use of nucleosome positioning *in situ* and cfDNA as a diagnostic marker. A) Nucleosome positioning acts as the cell memory at intermediate timescales between faster changes of reaction metabolites and long-term changes such as the accumulation of mutations and changes of DNA methylation. B) cfDNA extracted from blood plasma or other body liquids reflects the nucleosome positioning landscape in the cells of origin. This is because enzymes that shred chromatin into pieces in processes such as apoptosis, necrosis or NETosis preferentially cut DNA between nucleosomes.

Medical tests based on cfDNA are sometimes called “liquid biopsy” because this promising approach allows avoiding tissue biopsy in the case of solid tumours (Volik et al. 2016; Wan et al. 2017; Peng et al. 2020; Ignatiadis et al. 2021; Lo et al. 2021). The history of cfDNA research can be traced back to 1944 when it was first reported (Mandel and Metais 1948). cfDNA source was correctly interpreted as the products of apoptotic cleavage of chromatin subunits as early as 1970 (Williamson 1970; Henikoff and Church 2018). However, the active use of cfDNA for medical purposes using next generation sequencing (NGS) started only recently (Ignatiadis et al. 2021) with many diverse applications ranging from prenatal testing (Kitzman et al. 2012; Sun et al. 2018), cancer (Frenel et al. 2015; Phallen et al. 2017; Cristiano et al. 2019; Zviran et al. 2020), ageing (Teo et al. 2019), inference of patterns of gene expression (Snyder et al. 2016; Ulz et al. 2016) and transcription factor binding (Ulz et al. 2019), to even monitoring astronaut’s health on spaceflights (Bezdan et al. 2020). While the field of liquid biopsies is expanding dramatically, it is still in the search of methods balancing sensitivity and cost (Abbosh et al. 2017; Wan et al. 2019b; Peng et al. 2020).

Historically, the first class of genomics-based cfDNA diagnostic methods relied on mutation analysis (Frenel et al. 2015; Abbosh et al. 2017; Dudley and Diehn 2020; Zviran et al. 2020). Related approaches involve analyses of gene fusions (Palande et al. 2020) or copy number variations (CNVs) (Mouliere et al. 2018b). In all these cases, assay sensitivity critically depends on the sequencing depth as well as on the abundance of cfDNA derived from tumour cells (ctDNA) which usually correlates with the severity/stage of disease (Abbosh et al. 2017; van der Pol and Mouliere 2019; Zviran et al. 2020). In fact, a recent report showed that elevated cfDNA levels correlate with all-causes mortality (Kananen et al. 2020). Thus, many assays use cfDNA concentration as a marker of disease severity without sequencing.

However, if the detection method is based on few genomic regions that are not represented in cfDNA, then even increasing the sequencing depth would not help the diagnostics. To overcome this problem, it is possible to base cfDNA analysis on a larger number of genomic regions with more subtle epigenetic changes, hence, departing from the idea of mutation analysis and focusing the analysis on changes in DNA methylation (Shen et al. 2018; Erger et al. 2020; Liu et al. 2020; Nassiri et al. 2020) or hydroxymethylation (Song et al. 2017) of multiple genomic locations that reflect disease-specific changes in the cells of origin. cfDNA methylomics is being actively used in a growing number of applications. The main challenge with this class of approaches is that the detection of DNA modifications requires at least moderate sequencing depth which drives up the cost of the assay. In addition, changes in DNA modifications (as well as DNA sequence) accumulate at a long-term timescale and may not be prevalent at the onset of disease or as a response to therapy (see Figure 1A). To address these problems, one can consider assays that are based on the detection of smaller changes at a larger number of genomic loci. The most straightforward solution is to look at nucleosome positioning *per se* which is reflected in cfDNA localisation patterns.

New types of liquid biopsy tests based on nucleosome positioning-inspired analysis of cfDNA are sometimes termed “fragmentomics” and “nucleosomics” (Im et al. 2020). Fragmentomics analyses have been focused on the distribution of sizes of cfDNA fragments (Snyder et al. 2016; Underhill et al. 2016; Mouliere et al. 2018a; Sun et al. 2018; Markus et al. 2019; Guo et al. 2020; Zukowski et al. 2020) as well as the nucleotide patterns at their cut sites (Chandrananda et al. 2015). Sizes of cfDNA fragments reflect the contributions of different biological processes such as apoptosis, necrosis and NETosis. For example, apoptotic enzymes tend to cut out DNA fragments which are slightly smaller than mononucleosomal DNA (Serpas et al. 2019; Han et al. 2020). Such short cfDNA fragments tend to be enriched in cancer patients (van der Pol and Mouliere 2019). On the other hand, ultra-long cfDNA fragments may result from NETosis – a process in which neutrophils release nets of chromatin called neutrophil extracellular traps (NETs) in order to catch and destroy pathogens (Kustanovich et al. 2019). Such long cfDNA fragments can be associated with NETosis in different types of inflammation, for example, in diabetes (Wong et al. 2015) and COVID-19 (Ng et al. 2020). Necrotic cell death is also usually associated with longer DNA fragments (>10kb) (Kustanovich et al. 2019). Thus, each type of cell death has its distinct pattern of cfDNA size distribution. cfDNA size may also differ for different body fluids, e.g. urine usually harbours shorter cfDNA than blood plasma (van der Pol and Mouliere 2019). The situation is further complicated by the fact that cell senescence opposes cfDNA release (Rostami et al. 2020). Several studies in fragmentomics suggested using a simple ratio of the amount of short/long cfDNA fragments as an estimate of ctDNA/cfDNA fraction (Mouliere et al. 2018a; van der Pol and Mouliere 2019) but, given the complexity of different cell death pathways mentioned above, it is not always easily interpretable. We will show below that even within a narrow group of medical conditions the distribution of cfDNA sizes is quite heterogeneous. Another type of fragmentomics analysis is based on the fact that DNA nucleases have different sequence preferences (Serpas et al. 2019; Han et al. 2020) and therefore the distribution of nucleotide patterns at the ends of the cfDNA fragments may provide valuable diagnostic information (van der Pol and Mouliere 2019).

cfDNA nucleosomics is very promising since it eliminates the need of specific genomic markers and pre-set hypotheses about the underlying medical condition, and the bottleneck is now on the computational side. Recent studies have used machine learning to distinguish the cells of origin or perform binary classification healthy/cancer based on cfDNA density in promoters (Snyder et al. 2016; Wan et al. 2019a) or megabase-size genomic windows (Cristiano et al. 2019). Another successful approach combined several “simple” features in the PCA analysis including the amplitude of cfDNA oscillations with 10-bp periodicity, gene copy number variation and the relative abundances of cfDNA fragments with sizes in certain ranges (Mouliere et al. 2018a). One of the directions actively pursued by the cfDNA community is creating targeted sequencing assays based on nucleosomics of a small number of genomic regions – as small as just 6 regions in a recent publication (Zhu et al. 2021). The smaller the number of regions in the targeted nucleosomics assays the better, but this also has to be balanced with the sensitivity and ability to recognise more than one medical condition. Currently, the “holy grail” of liquid biopsies – the ability to diagnose an arbitrary medical condition – is still far from reach. Notably, achieving this aim requires access to as many as possible published cfDNA datasets to train the models. Few web sites started appearing that allow visualisation and download of a limited number of cfDNA datasets (Yu et al. 2020; Zheng et al. 2020), but a centralised resource which collects cfDNA datasets from the dozens of currently available (and increasing in number) publications is currently not available. Here, we have developed such a resource - NucPosDB, which aims to curate all published datasets of sequenced cfDNA, nucleosome positioning maps *in vivo* and software for nucleosomics analysis. NucPosDB also intends to provide our integrative analysis to quantify the genome in terms of regions with differential nucleosome occupancy and stability (Vainshtein et al. 2017), providing a connection between nucleosome maps in healthy (Schones et al. 2008; Gaffney et al. 2012) and cancer human cells (Mallm et al. 2019).

## Construction and content

### Database structure

NucPosDB curates open- and restricted-access datasets of nucleosome positioning *in vivo* and sequenced cfDNA, as well as computational software for cfDNA/nucleosome positioning analysis and modelling. The structure of the database is summarised in Figure 2. It contains the following sections: (1) nucleosome positioning in vivo, (2) sequenced cfDNA, (3) database of regions in the human genome with stable nucleosome occupancy for a given condition, and the repository of software for nucleosomics, further separated into three subsections devoted to (4) analysis of nucleosome maps *in vivo*, (5) prediction of nucleosome formation preferences based on DNA sequence and (6) cfDNA-specific analysis.

**Figure 2.**
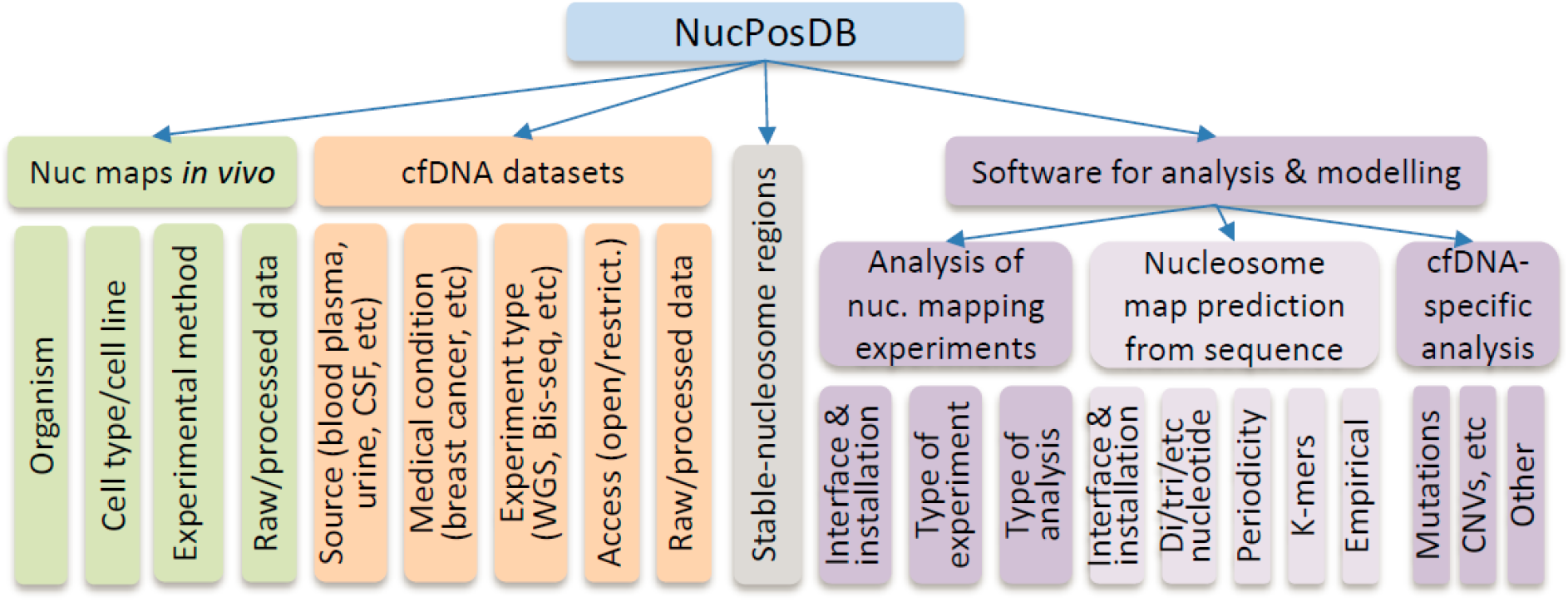
The structure of NucPosDB containing six major sections (listed left to right in the scheme): (1) nucleosome maps measured in vivo in different cell types, (2) sequenced cfDNA datasets, (3) regions with stable nucleosome occupancy in the human genome for different conditions based on (1) and (2), (4) software for analysis of nucleosome mapping experiments, (5) software for predicting preferences of nucleosome formation from the DNA sequence and (6) software for cfDNA-specific analysis.

The section of nucleosome positioning *in vivo* contains datasets from >250 publications in >16 biological species, dominated by *Saccharomyces cerevisiae* (28.6%), *Mus musculus* (25.9%), *Homo sapiens* (20.1%) and *Drosophila melanogaster* (14.3%). Figure 3 demonstrates relative abundances of different model organisms used for nucleosome positioning analysis. This section of the database features more than 18 experimental techniques, dominated by MNase-seq, complemented by methods such as histone H3 ChIP-seq, MH-seq, MPE-seq, MiSeq, NOME-seq and RED-seq (detailed in our previous publications (Teif 2016; Teif and Clarkson 2019) as well as newer techniques based on long single-molecule reads, Nanopore-seq (Baldi et al. 2018) and Fiber-seq (Stergachis et al. 2020), and nucleosome-scale mapping of 3D genome contact, Micro-C (Hsieh et al. 2015). Techniques such as ATAC-seq, which map nucleosomes only in a limited number of “open” genomic locations are currently not included in NucPosDB.

**Figure 3.**
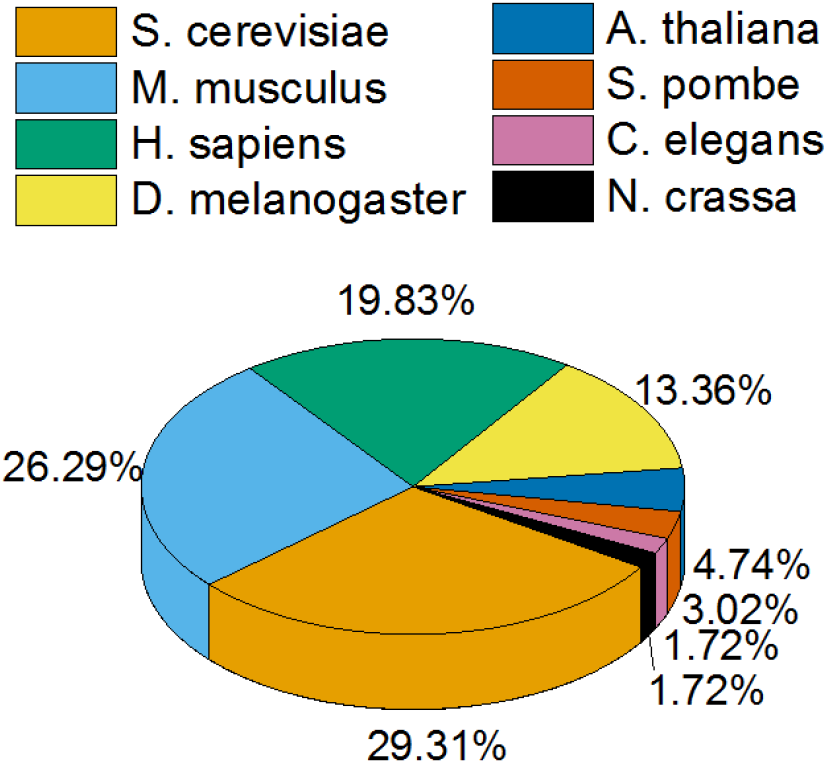
The distribution of nucleosome positioning datasets across different biological species.

The repository of sequenced cfDNA represents a recent addition to NucPosDB and currently features more than 75 studies. cfDNA processing is complicated by the fact that many datasets dealing with patient data have restricted access, e.g. where the raw data is stored in the European Nucleotide Archive (ENA) or the database of Genotypes and Phenotypes (dbGaP). The application for access to each such dataset is considered individually by the corresponding data access committee and the time required to receive regulatory approval may reach several months. On the other hand, when the raw data is stored in databases such as GEO, such datasets are available without restrictions. NucPosDB curates both open-access and restricted-assess datasets, but only open-access datasets are supplied with the processed data including the locations of all mapped nucleosomes and stable-nucleosome regions (see below). Table 1 shows examples of cfDNA datasets from NucPosDB that have no access restrictions. cfDNA datasets included in NucPosDB can be browsed by organism (e.g. human, mouse or dog). Currently, the majority of cfDNA datasets included in NucPosDB are of human origin. For patients, it is possible to select medical condition (currently around 50 conditions), source of cfDNA (blood e.g. serum/plasma, cerebrospinal liquid or urine), experimental method (currently 12 methods) and access type (restricted or not).

**Table 1.**
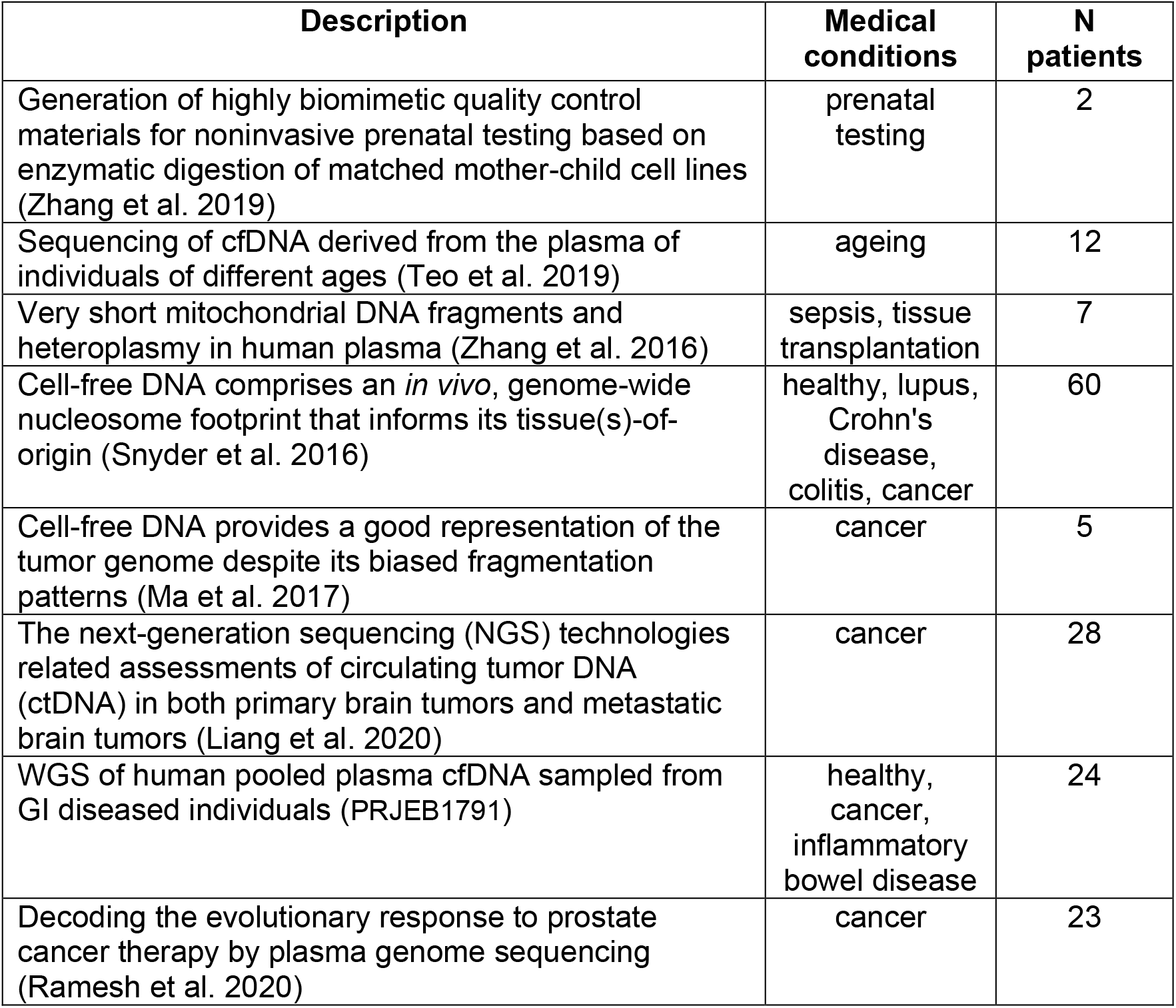
Example open-access datasets from NucPosDB reporting whole genome sequencing of cfDNA.

A special NucPosDB section is devoted to the stable-nucleosome regions of the human genome. It contains condition-specific coordinates of genomic locations where nucleosome occupancy has low relative standard deviation across all samples within the same condition. This is defined with NucTools (Vainshtein et al. 2017) using a window-based approach as detailed below and arranged in tab-separated BED files with the following columns: chromosome, region start, region end, normalised nucleosome occupancy, standard deviation, relative deviation. In addition, for a number of open-access cfDNA entries our database provides access to the uniformly processed BED files with locations of all mapped nucleosomes (based on paired-end cfDNA reads). We have mapped these cfDNA reads to the human genome assemblies hg19 and hg38 as detailed below. These were further processed with NucTools (Vainshtein et al. 2017) to generate tab-separated files with the following columns: chromosome, fragments start, fragment end, fragment size. Each patient’s sample has been processed separately. The links from the interactive database tables lead to the file repository with directories separated by publication and further split into different medical conditions.

The repository of software for analysis of nucleosome positioning experiments currently contains 31 entries representing different classes of software ranging from nucleosome array visualisers and nucleosome peak callers to predictors of specific parameters such as the nucleosome repeat length (Vainshtein et al. 2017). The repository of algorithms for prediction of DNA sequence-dependent affinity of nucleosome octamer currently contains 22 entries, as described previously (Teif 2016; Teif and Clarkson 2019). The repository of software specific for the analysis of cfDNA currently includes 32 entries.

### Data collection and curation

The datasets were searched in NCBI GEO as well as in peer-reviewed publications and preprints from bioRxiv and medRxiv servers. Initial search was conducted using the keywords “nucleosome positioning”, “MNase-seq” and “cfDNA”. Further relevant studies were extracted through publication chaining. Over 300 papers reporting relevant datasets and software were arranged into five sections: nucleosome maps *in vivo*, cfDNA datasets, computational tools for nucleosome positioning analysis and modelling, and cfDNA analysis. The criterion for the dataset inclusion was the ability to reconstruct based on a given dataset a nucleosome positioning profile with single-nucleotide resolution. Datasets reporting methods such as ChIP-seq and microarrays were normally excluded unless the corresponding publications provided specific nucleosome positioning analysis. ATAC-seq was excluded since it maps nucleosomes only in a limited number of “open” genomic locations. cfDNA datasets were included when they were obtained using any variation of a sequencing technique that involves whole-genome or targeted sequencing and thus allows partial or complete reconstruction of nucleosome profiles. This includes methods determining DNA methylation and hydroxymethylation, but not microarray-based techniques/assays.

### User interface

The user interface of NucPosDB is realised in PHP. The search and keywords selection is currently enabled with the help of the TablePress plugin for WordPress (tablepress.org, author Tobias Bäthge, Magdeburg, Germany). Datasets can be searched by typing a query into the search box or using multiple-item selection in drop-down menus such as “Organism” and “Experiment type”. Additionally, the repository of cfDNA datasets contains drop-down menus “Medical condition”, “cfDNA source” and “Access” (open/restricted). The interactive tables with experimental datasets can be ordered or sub-selected by the combination of the following criteria: “Description” (typically includes the title of the original publication and a clickable link), “Organism”, “Cell type” (only in the section nucleosome maps *in vivo*), “Experiment type/method”, “Raw data” and “Processed data”. The cfDNA repository allows additional selection/ordering criteria: “Medical condition”, “cfDNA source”, “Access” (open/restricted) and “Number of patients”.

### Data processing

The calculation of the histogram of DNA fragment size distribution was carried out using R. The calculation of nucleotide frequencies was performed with HOMER (Heinz et al. 2010). Raw paired-end reads were aligned to the human genomes hg19 and hg38 using Bowtie (Langmead et al. 2009), reporting only uniquely aligned reads with up to two mismatches. Normalised nucleosome occupancy was calculated genome-wide with 100-bp windows by dividing the average nucleosome occupancy in a given window by the average chromosome-wide nucleosome occupancy. Stable-nucleosome regions were determined with NucTools with 100 bp sliding window and the threshold 0.5 applied to the relative deviation of nucleosome occupancy across all samples with a given condition, as described previously (Vainshtein et al. 2017). The relative deviation was defined as the ratio of the standard deviation to the normalised nucleosome occupancy in a given window.

## Utility and discussion

One of the main purposes of having a centralised repository of nucleosome positioning/cfDNA datasets is to be able to assess the data heterogeneity within conditions and the variability between different conditions and experimental protocols. While a systematic analysis of such variability of all datasets in NucPosDB is beyond the scope of the current work, let us demonstrate here typical distributions of two basic characteristics of cfDNA, namely the GC content and the DNA fragment sizes.

Firstly, let us consider the nucleotide frequency as a function of the distance from the cfDNA fragment end (Figure 4A). This type of analysis is motivated by previous findings that endogenous nucleases have distinct preferences for DNA cut sites, and these preferences are different from artificial cut sites observed in MNase-seq experiments (Serpas et al. 2019; van der Pol and Mouliere 2019; Han et al. 2020). Apoptosis in different types of cancer may involve the same set of nucleases, therefore, based on this metric, different types of cancer may not be easily distinguishable from each other. Indeed, this is what we observe for the distribution of GC frequencies near cfDNA fragment ends in Figure 4A. On the other hand, different biological processes such as NETosis may employ a different combination of enzymes, thus it may be possible to distinguish medical conditions that are characterised by inflammation (inflammation triggers NETosis). Indeed, Figure 4A shows that nucleotide profiles of cfDNA from patients with lupus (systemic inflammation) differ quite significantly from those in cancer or healthy controls.

**Figure 4.**
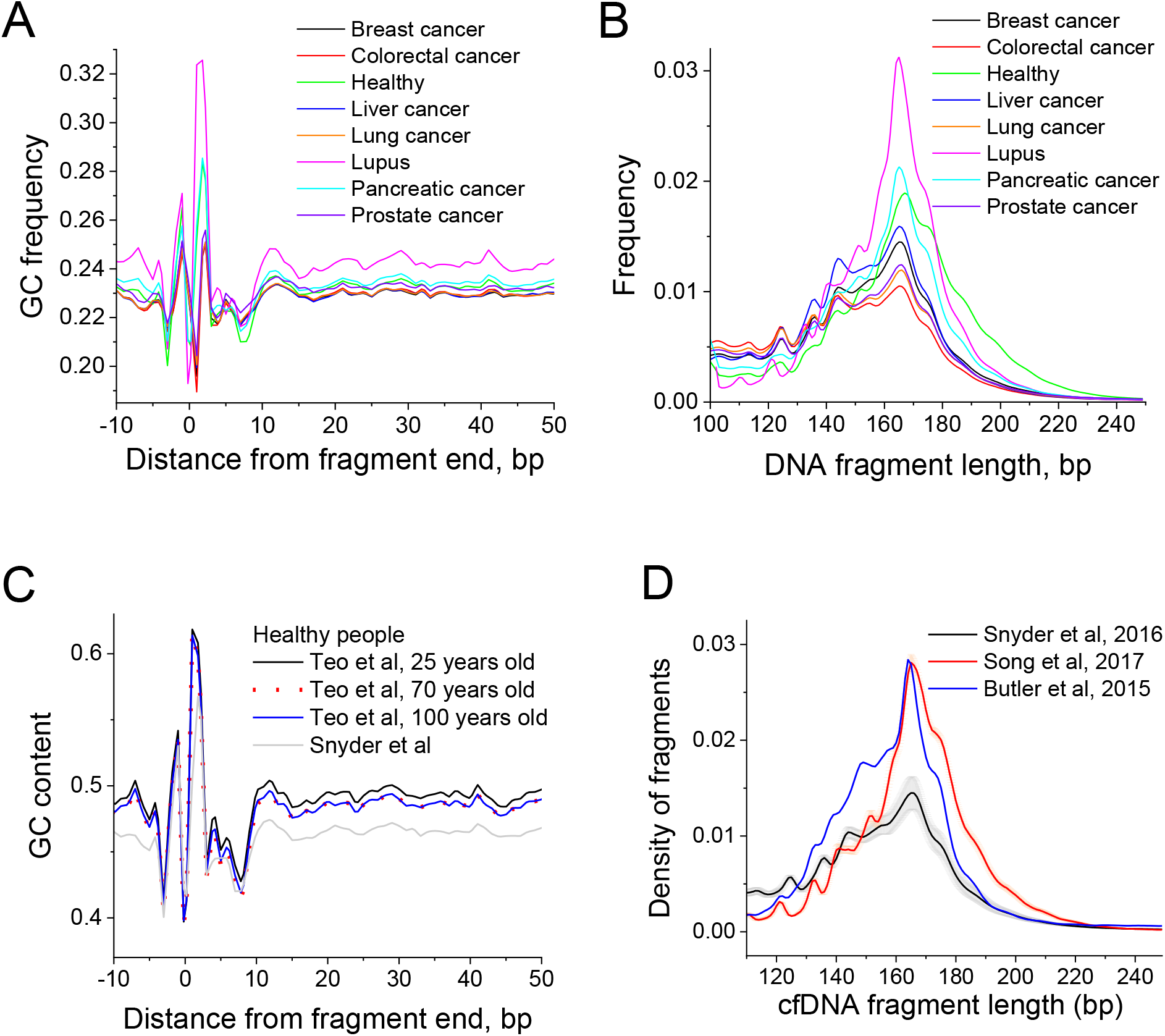
Aggregate characteristics of cfDNA datasets across different medical conditions (A-B) and ages of healthy people (C-D). A) GC content as a function of the distance from the end of cfDNA fragment (Snyder et al. 2016). B) Distribution of lengths of cfDNA fragments (Snyder et al. 2016). C) GC content as a function of the distance from the end of cfDNA fragment for 25, 70 and 100-years old people (Teo et al. 2019), compared with pooled healthy people from another study (Snyder et al. 2016). D) Differences of cfDNA fragment sizes for cfDNA of breast cancer patients collected in three different studies (Snyder et al. 2016, Song et al., 2017 and Butler et al, 2015).

Next, let us consider the distributions of DNA fragment sizes. Previous studies reported that cancer cfDNA appears to have shorter fragments that are more strongly digested (Snyder et al. 2016; Underhill et al. 2016; Mouliere et al. 2018a; Sun et al. 2018; Markus et al. 2019; Guo et al. 2020; Zukowski et al. 2020). Our results do show differences in cfDNA fragment size distributions, most notably for lupus (Figure 4B). The difference of cfDNA in lupus from cancer and healthy samples may be explained by the different DNA digestion processes undergoing in this systemic inflammatory condition (Figure 1B). However, special care is required to normalise the data and take into account different protocols (e.g. the lupus samples in the study considered above may have been clinically processed in a different way than the cancer samples).

The differences in the experimental protocols used in different labs for cfDNA processing as well as comorbidities of patients may play major roles in the data interpretation. To demonstrate this, Figure 4C and D compare samples from different subgroups of healthy people. Figure 4C shows that the average GC content of cfDNA extracted for whole-genome sequencing by different methods differs dramatically. In one case, Teo et al collected cfDNA from three age groups (25, 70 and 100 years old) and the differences of GC profiles between these age groups are pretty minor (Teo et al. 2019). On the other hand, in another group of healthy people where sequencing was performed by the method of Snyder et al the average GC content is about 3% lower (Snyder et al. 2016). Such difference may lead to biased representation of different types of genomic regions and needs to be taken into account when comparing datasets across different laboratories. Indeed, Figure 4D shows that the distribution of cfDNA fragment sizes varies quite substantially between datasets reported by three different labs even when all of these refer to the same condition (breast cancer in this example), and when samples within one lab’s dataset are consistently similar to each other. This probably reflects differences in experimental protocols and needs to be taken with special care when performing nucleosomics analysis for cancer diagnostics. Similar care is needed when comparing MNase-seq datasets obtained in different laboratories, because it is known that parameters such as the degree of chromatin digestion greatly affect nucleosome maps due to differential sensitivity of partially unwrapped nucleosomes to digestion level (Teif et al. 2014; Chereji et al. 2016; Ramachandran et al. 2017). In such situations it may be helpful to adjust clinically relevant analyses taking into account the locations of regions with stable nucleosome occupancy in a given condition as reported by NucPosDB.

Finally, the examples shown above demonstrate that the development of a robust clinical diagnostics based on cfDNA nucleosomics will require many datasets across different laboratories and types of wet lab assays. This is where NucPosDB may be particularly helpful, allowing the use the data from more than 10,000 patients.

## Conclusions

NucPosDB offers a user-friendly interface and curates published *in vivo* nucleosome positioning datasets including >18 types of experimental techniques in >16 different species and distinct cell types, supplemented with the repository curating cfDNA datasets for more than 10,000 patients as well as the software packages for “nucleosomics” analysis. For many open-access datasets we also provide systematically calculated condition-specific stable-nucleosome regions which are useful in comparison between different conditions. In the future, NucPosDB can serve as a centralised resource for the nucleosomics community, providing a platform for the annotation of cfDNA datasets and storage of processed data required for training models for patient diagnostics with liquid biopsies.

## Declarations

### Availability of data and materials

NucPosDB is available at https://generegulation.org/nucposdb/.

### Authors’ contributions

MS developed the repository of cfDNA, curated the repository of nucleosome positioning *in vivo* and performed cfDNA data analysis. KVP contributed to the curation of the repository of nucleosome positioning *in vivo* and data analysis. SPA compiled the repository of software for cfDNA analysis and contributed to cfDNA analysis. DRJ contributed to cfDNA analysis. VBT designed and supervised the project, performed programming and drafted the manuscript.

